# Daily running enhances molecular and physiological circadian rhythms in skeletal muscle

**DOI:** 10.1101/2020.10.19.346015

**Authors:** Nuria Casanova-Vallve, Drew Duglan, Megan E. Vaughan, Michal K. Handzlik, Weiwei Fan, Ruth T. Yu, Christopher Liddle, Michael Downes, Julien Delezie, Rebecca Mello, Alanna B. Chan, Marie Pariollaud, Pål O. Westermark, Christian M. Metallo, Ronald M. Evans, Katja A. Lamia

## Abstract

**Objective:** Exercise is a critical component of a healthy lifestyle and a key strategy for the prevention and management of metabolic disease. Identifying molecular mechanisms underlying adaptation in response to chronic physical activity is of critical interest in metabolic physiology. Circadian rhythms broadly modulate metabolism, including muscle substrate utilization and exercise capacity. Here, we define the molecular and physiological changes induced across the daily cycle by voluntary low intensity daily exercise.

**Methods:** Wildtype c57BL6/J male and female mice were housed with or without access to a running wheel for six weeks. Maximum running speed was measured at four different zeitgeber times (ZTs, hours after lights on) using either electrical or manual stimulation to motivate continued running on a motorized treadmill. RNA isolated from plantaris muscles at six ZTs was sequenced to establish the impact of daily activity on genome-wide transcription. Patterns of gene expression were analyzed using Gene Set Enrichment Analysis (GSEA) and Detection of Differential Rhythmicity (DODR). Blood glucose, lactate, and ketones, and muscle and liver glycogen were measured before and after exercise.

**Results:** We demonstrate that the use of mild electrical shocks to motivate running negatively impacts maximum running speed and describe a manual method to motivate running in rodent exercise studies. Using this method, we show that time of day influences the increase in exercise capacity afforded by six weeks of voluntary wheel running: when maximum running speed is measured at the beginning of the nighttime active period in mice, there is no measurable benefit from a history of daily voluntary running, while maximum increase in performance occurs at the end of the night. We show that daily voluntary exercise dramatically remodels the muscle circadian transcriptome. Finally, we describe daily rhythms in carbohydrate metabolism associated with the time-dependent response to moderate daily exercise.

**Conclusions:** Collectively, these data indicate that chronic nighttime physical activity dramatically remodels daily rhythms of muscle gene expression, which in turn support daily fluctuations in exercise performance.

## 1. Introduction

Exercise performance can be modulated by diet, metabolism, and environmental factors including the time of day [1]. In athletes, physical performance is enhanced in the evening compared to the same exercise in the morning [2–5]. Similarly, sedentary mice exhibit improved endurance capacity late in the active period [6]. Thus, in both humans and rodents, time of day influences maximum exercise capacity. Further, it has recently become clear that the time at which exercise is performed influences its impacts on physiology. Notably, the time of day at which exercise is performed alters its beneficial impact in metabolically compromised patients [7; 8]. Taken together, these findings indicate that exercise physiology and circadian rhythms are intimately intertwined and motivate our search for a deeper understanding of how they are connected at the molecular level.

The circadian clock is based on transcriptional feedback loops [9], in which brain and muscle ARNT-like protein 1 (BMAL1) and circadian locomotor output cycles kaput (CLOCK) regulate thousands of target genes, including their repressors encoded by the period (*Per1, Per2*, and *Per3*) and cryptochrome (*Cry1* and *Cry2*) genes. Circadian clocks dictate the rhythmic expression of metabolic target genes in anticipation of predictable daily fluctuations in demand. In skeletal muscle, genes involved in carbohydrate and lipid metabolism are enriched among those that exhibit rhythmic expression and those that are regulated by BMAL1 [10; 11]. Consistent with these findings, the metabolic state of muscles and their response to acute exercise depends on time of day [12; 13]. Moreover, deletion of *Bmal1* results in impaired glucose metabolism [14; 15], reduced grip strength, and increased muscle fibrosis [16]. Deletion of the circadian repressors *Cry1* and *Cry2* results in enhanced exercise performance and metabolic regulation with increased fatty acid oxidation, involving activation of PPARδ [17].

The potential for circadian rhythms to influence exercise outcomes has long been recognized both in real world and laboratory settings; time of day was included among the parameters to be considered in the design of exercise experiments involving animals by an expert panel of the American Physiological Society as early as 2006 [18]. That monograph also included a discussion of the use of “aversive stimuli” to motivate laboratory animals to exercise. The authors urged careful consideration of the use of mild electric shocks to motivate animals to run, mostly due to ethical considerations. Although electrical stimulation via the neuromuscular junction is well established to be a key physiological stimulant of muscle contraction, and electrical stimulation of muscle fibers *in vitro* is widely used to study physiological responses [19–21], surprisingly little research has addressed the potential confounding effect of the use of mild electrical stimulation in rodent exercise testing.

Here, we demonstrate that the impact of daily moderate exercise on maximum sprint speed depends on the time of day. Notably, we find that the widespread use of electrical stimulation to motivate running in rodent exercise testing dramatically alters experimental outcomes. We provide a comprehensive characterization of daily transcriptome remodeling in response to running wheel (RW) access that reveals specific alterations in carbohydrate and lipid metabolism that support time of day differences in high- and low-impact exercise. We show that glycogen and lipid metabolism connect circadian rhythms and exercise physiology and establish resources that will enable further investigation of the molecular mechanisms underlying circadian rhythms in optimal exercise performance and daytime-dependent responses to physical activity.

## 2. Materials and Methods

### 2.1 Mouse Models

C57BL/6J mice were purchased from the Scripps Research breeding colony at six weeks of age. They were maintained in standard 12:12 light:dark conditions unless otherwise indicated and were group housed except when given voluntary access to running wheels, in which case they were singly housed in running wheel cages. They were given ad libitum access to normal mouse chow and water. All animal care and treatments were approved and overseen by The Scripps Research Institutional Animal Care and Use Committee under protocol #10-0019, comply with the ARRIVE guidelines and were carried out in accordance with the National Research Council’s Guide for the Care and Use of Laboratory Animals.

### 2.2 Treadmill Exercise Testing

Treadmill running performance was assessed in 5-12 mice per group. Untrained, sedentary male mice were tested at 9 weeks of age, and then underwent six or eight weeks of voluntary running wheel training and were tested again within 30 minutes of being removed from running wheel cages. Blood glucose (Aviva Accu-chek, Roche Diagnostics), lactate (Lactate Scout, SensLab) and ketone (Precision Xtra, Abbott Laboratories) levels were assessed via the tail vein at baseline (week prior to experimentation) and immediately post-sprint run.

### 2.3 RNA extraction, quantitative PCR, and RNA sequencing

RNA was extracted from quadriceps or plantaris muscle tissue by homogenization in QIAzol (Qiagen) or Trizol (Invitrogen) by standard methods. For RNA sequencing, total mouse plantaris RNA was isolated using Trizol (Invitrogen) and the RNeasy mini kit with on-column DNase digestion (Qiagen). Sequencing libraries were prepared from 100 ng of total RNA using the TruSeq Stranded Total RNA sample preparation kit (Illumina) according to the manufacturer’s protocol. Libraries prepared from three biological replicates for each experimental timepoint were subjected to single-ended sequencing on the Illumina HiSeq 2500 using barcoded multiplexing and a 90-bp read length. Differential gene expression analysis, statistical testing and annotation were performed using Cuffdiff v2.2.1 [22]. Transcript expression was calculated as gene-level relative abundance in fragments per kilobase of exon model per million mapped fragments (FPKM) and employed correction for transcript abundance bias [23].

### 2.4 Tissue glycogen analysis

Quadriceps and liver glycogen content was assessed from tissues dissected under basal conditions or immediately post-sprint exercise testing. Glycogen was detected using a commercial glycogen assay kit (Sigma, MAK016), as per manufacturer’s instructions, including a step to subtract the background free glucose present before enzymatic digestion of glycogen. Tissue (~50-100 mg) was homogenized in MQ water and glycogen concentration determined by a coupled enzymatic assay. The colorimetric reaction was detected at 570 nm in a microplate reader (VersaMax).

### 2.5 Serum triglyceride analysis

Circulating triglycerides were assessed immediately post-sprint exercise testing. Blood was collected by cardiac puncture and left undisturbed at room temperature for 30 min. After centrifugation at 4°C for 10 min at 2000g, the serum supernatant was collected. Triglycerides were detected from serum using a commercially available triglyceride assay kit (Sigma, MAK266). Triglyceride concentration was determined by enzymatic breakdown to glycerol. The colorimetric reaction from glycerol oxidation was detected at 570 nm in a microplate reader (VersaMax).

### 2.6 Serum NEFA analysis

Circulating non-esterified fatty acids (NEFAs) were assessed immediately post-sprint exercise testing. Serum was isolated as described above. NEFAs were detected from serum using a commercially available NEFA assay reagents (Wako Diagnostics, 999-34691). NEFA concentration was determined by the enzymatic acylation of coenzyme A; the colorimetric reaction from Acyl-coA oxidation was detected at 550 nm in a microplate reader (VersaMax).

### 2.7 Tissue polar metabolite extraction

Frozen tissue samples (20-30 mg) were homogenized for 2 min using ceramic beads (Precellys 2 mL Hard Tissue Homogenizing Ceramic Beads Kit, Bertin Instruments, US) in 500 μL −20 °C methanol, ice-cold 400μL saline and ice-cold dH_2_O containing amino acid isotope labelled internal standards (Cambridge Isotope Laboratories, #MSK-A2-1.2). An aliquot of homogenate (50μL) was dried under air and resuspended in RIPA buffer for protein quantification using BCA assay (BCA Protein Assay, Lambda, Biotech Inc., US). To the remaining homogenate, 1 mL of chloroform was added and the samples were vortexed for 5 min followed by centrifugation at 4°C for 5 min @ 15 000g. The organic phase was collected and the remaining polar phase was re-extracted with 1 mL of chloroform. An aliquot of the polar phase was collected and vacuum-dried at 4°C and subsequently derivatized with 2% (w/v) methoxyamine hydrochloride (Thermo Scientific) in pyridine for 60 min following by 30 min sialyation N-tertbutyldimethylsilyl-N-methyltrifluoroacetamide (MTBSTFA) with 1% tertbutyldimethylchlorosilane (tBDMS) (Regis Technologies) at 37°C. Polar derivatives were analyzed by GC-MS using a DB-35MS column (30m x 0.25mm i.d. x 0.25 μm, Agilent J&W Scientific) installed in an Agilent 7890A gas chromatograph(GC) interfaced with an Agilent 5975C mass spectrometer (MS) as previously described [24].

### 2.8 Tissue triglyceride extraction

Frozen tissue samples (20-30 mg) were homogenized for 2 min using ceramic beads (Precellys 2 mL Hard Tissue Homogenizing Ceramic Beads Kit, Bertin Instruments, US) in 500 μL −20 °C methanol, ice-cold 400μL saline and ice-cold dH_2_O containing C12 MAG (1-0-dodecyl-rac-glycerol, #SC-201977, Santa Cruz) as internal standard. To the remaining homogenate, 1 mL of chloroform was added and the samples were vortexed for 5 min followed by centrifugation at 4°C for 5 min @ 15 000g. The organic phase was collected and 2 μL of formic acid was added to the remaining polar phase, which was re-extracted with 1 mL of chloroform. Combined organic phases were dried under air flow and the pellet was resuspended in 125 μL of chloroform. Triglyceride species were separated chromatographically on a C5 column (Luna, 100 x 2 mm, 5 μm, Phenomenex). Mobile phase A was composed of 95:5 ratio of water:methanol, 0.1 % formic acid and 5 mM ammonium formate, and mobile phase B was 60:35:5 isopropanol:methanol:water, 0.1% formic acid and 5 mM ammonium formate. The chromatography gradient elution was as follows: 0 min, 0% B, flow rate 0.1 ml/min; 5 min, 0% B, flow rate 0.4 ml/min; 50 min, 100% B, flow rate 0.4 ml/min; 50.1 min, 100% B, flow rate 0.5 ml/min, 67 min; 100% B, flow rate 0.5 ml/min; 67.5 min, 0% B, flow rate 0.5 ml/min and 76 min, 0% B, flow rate 0.5 ml/min. Liquid chromatography mass spectrometry was performed on an Agilent 6460 QQQ LC-MS/MS instrument. Triglyceride species were analyzed by SRM of the transition from precursor to product ions at associated optimized collision energies and fragmentor voltage.

### 2.9 Statistical analysis

Statistical analyses were performed using GraphPad Prism software. Unless otherwise indicated, ANOVA was used to determine significance with a threshold of 0.05 acceptable false positive (P < 0.05).

## 3. Results

### 3.1 Electrical stimulation can alter outcomes in rodent treadmill tests

We abandoned the common practice of using aversive electrical stimulation several years ago in favor of using gentler physical stimuli to motivate continued running because we suspected that even mild electric shocks may influence outcomes [17], a view that is informally shared by many in the exercise physiology community. To directly evaluate this idea, we compared maximum sprint speeds measured when continued running was motivated by either mild electrical stimulation or by our manual method. Indeed, the use of electrical stimulation in exercise testing profoundly impairs outcomes (Figure 1A and Figure S1A).

**Figure 1:**
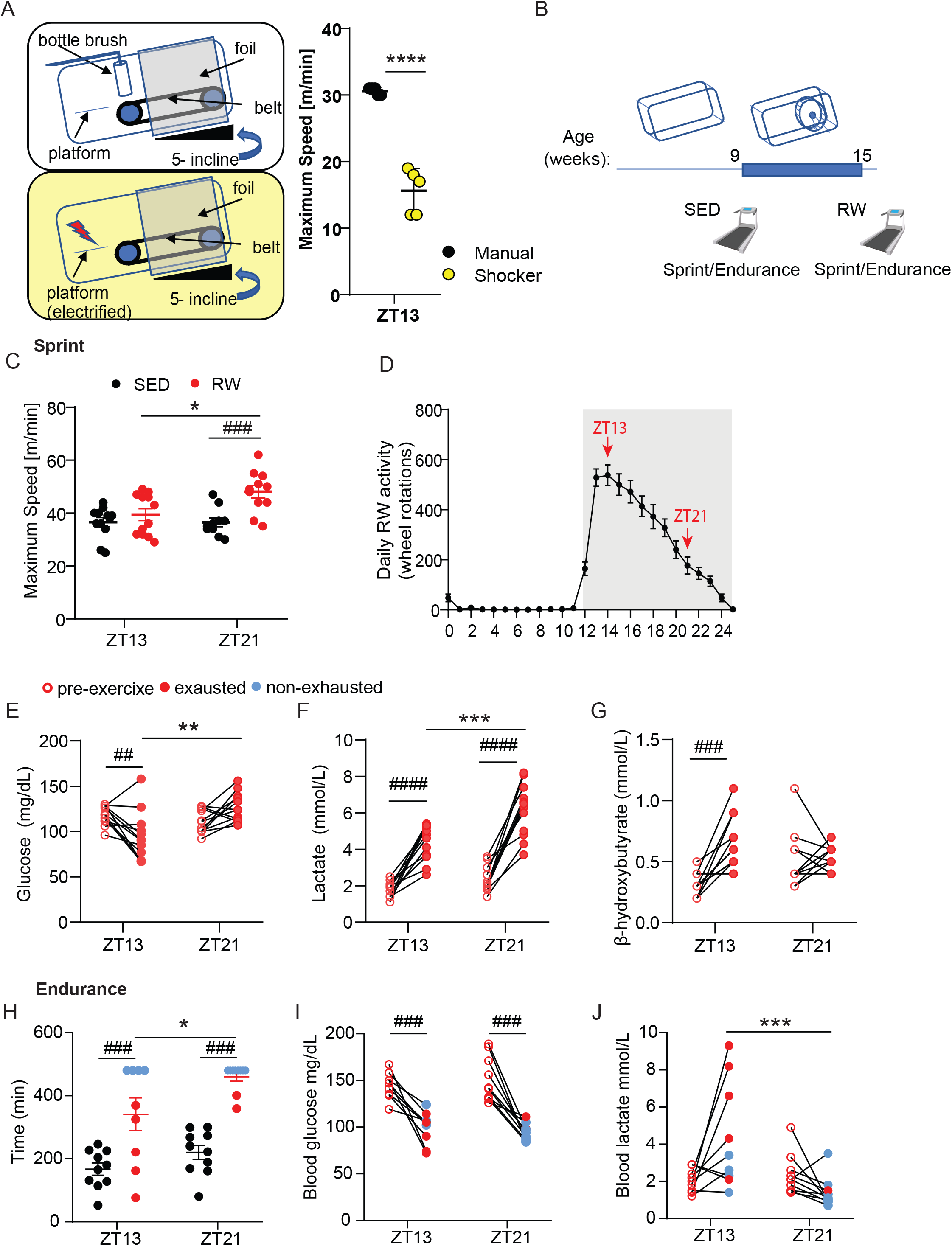
Time of day impacts maximum speed and endurance exercise in trained but not sedentary mice. (A) Diagram of manual and electrical aversive stimuli used to motivate continued treadmill running and maximum speeds achieved at ZT13 by c57BL/6J male mice using manual (black) or electrical (yellow) stimuli. (B) Schematic diagram of the experimental setup. When c57BL/6J mice were 9 weeks old, their maximum sprint speed or running endurance was determined using a motorized treadmill at either zeitgeber time (ZT, hours after lights on) 13 or 21. Then, they were transferred to cages with running wheels for six weeks, and their speed and endurance were determined at either ZT13 or ZT21 again. (C,H) Maximum speeds (*C*) or running time (*H*) achieved by male mice described in (*B*). (D) Temporal profile of running wheel activity during the training period, in one-hour bins. For each ZT, the total activity recorded in the preceding hour is shown. Data represent the mean ± s.e.m. for six mice per group. (E-G,I,J) Blood glucose (E,I), lactate (F,J), and beta-hydroxybutyrate (G) before (open circles) or after (filled circles) sprint (E-G) or endurance (I,J) test in mice housed with running wheels for six weeks. In (*C,H*) each black or red symbol represents a unique sedentary or active mouse, respectively. * P < 0.05, ** P < 0.01 *** P < 0.001 by two-way ANOVA followed by Tukey’s multiple comparison test. In (*E,F,G,I,J*) each connected pair of open and filled circles represents measurements from the same individual mouse before (open circles) and after (closed circles) the exercise test. *** P < 0.001 by mixed-effects analysis with Sidak’s multiple comparison test. #### P < 0.0001 by paired t-test.

### 3.2 Daily voluntary running increases maximum speed at ZT21 but not at ZT13 in male mice

Given the significant impact of genetically deleting both of the circadian repressors *Cry1* and *Cry2* on maximum treadmill running speed in mice [17], we hypothesized that maximum sprint speed may depend on the time of day. Others have reported that sedentary male mice exhibit enhanced sprint exercise capacity early in the night when CRY1/2 protein expression is lower [6] or at the dark to light transition [25]. However, we did not detect any significant impact of time of day on maximum sprint speed in sedentary c57BL/6J male mice (Figures 1B, 1C, and S1B). Notably, maximum speeds reported in a similar study [6] (38-42 cm/s, equivalent to 22-25 m/min) were much slower than the speeds that we measured (30-35 m/min, equivalent to 50-58 cm/s). The only major difference in the testing protocols is the aversive stimulus used to motivate mice to continue running.

Given the importance of metabolism for driving peripheral circadian rhythms as exemplified by the role of feeding time in setting the timing of peripheral clocks [26], we hypothesized that daily exercise would enhance daily rhythms in muscles, and may reveal an impact of time of day on maximum running speed. After six weeks of voluntary training by running wheel access, the enhanced speed afforded by wheel running activity is significantly greater during the nighttime active period (Figures 1B, 1C, and S1B) when mice are given continuous free access to a running wheel until the treadmill test is performed. Surprisingly, there is no measurable increase in maximum sprint speed after six weeks of running wheel training when sprint testing is performed at zeitgeber time (ZT, hours after lights on) ZT13 (early dark phase which is the active period for nocturnal rodents, equivalent to morning in humans). The increased sprint capacity after voluntary training when measured at ZT6 (Figure S1B) is similar to what we have measured in the past at ZT8 [17], demonstrating the consistency of our measurements across several years and when the assay is performed by different individual investigators. Importantly, the increased speed is due to voluntary running wheel activity rather than age as male mice maintained in a cage without running wheel access did not improve their performance (Figure S1C). As expected, mice with access to running wheels gain less weight than sedentary counterparts (Figure S1D). Notably, the temporal pattern of voluntary running wheel activity does not match the temporal pattern of maximum sprint capacity, as voluntary running at ZT13 is greater than at ZT21 (Figure 1D), while maximum trained speed is greater at ZT21. Finally, the impact of time of day on maximum sprint exercise capacity after six weeks of voluntary training was only observed in male mice. Female mice tend to be more active than males when provided running wheels (Figures S1E). Trained female mice have greater maximum sprint speed than their male counterparts, which was not different between ZT13 and ZT21 (Figure S1F).

To investigate mechanism(s) underlying the observed preferential increase in maximum speed in the late active phase after six weeks of running wheel access, we measured circulating metabolites before and after exercise testing at ZT13 and ZT21. Male mice maintain stable levels of glucose and beta hydroxybutyrate after the sprint test at ZT21, while at ZT13, they experience a significant reduction of blood glucose and increase in circulating ketone bodies (Figure 1E-G). Blood lactate increased after exercise at both times of the day, with a greater elevation at ZT21 (Figure 1F). Differences in circulating metabolites after exercise could reflect differences in the amount of wheel running preceding the exercise tests (Figure 1D). However, the impact of ZT on the responses of blood lactate and glucose to an acute exercise stimulus was similar in sedentary mice, in which ZT did not affect exercise capacity (Figure S1G,H). These data suggest that enhanced maximum speed at ZT21 in mice that experience daily moderate activity is associated with increased selective utilization of carbohydrates for energy production during exercise at ZT21. Consistent with this proposition, a recent study showed that humans exhibit a significantly increased respiratory exchange ratio (RER, a measure of the relative utilization of carbohydrates and fats for energy production) when performing exercise in the afternoon compared to the same exercise performed in the morning [27]. Similarly, the impact of exercise on lipids, carbohydrates, and other metabolites in several mouse organs depends on the time of day at which exercise is performed [28]. However, these differences in substrate utilization seem to be insufficient to increase maximum speed at ZT21 since they are also observed in sedentary animals, which have similar maximum running speeds at ZT13 and ZT21.

As reported by others [6], we find that sedentary mice tend to have greater treadmill running endurance at ZT21 than at ZT13 (Figure 1H). Voluntary access to the RW increased running endurance as expected. Indeed, the endurance of mice housed with running wheels is so great that most of the animals tested at ZT21 fail to reach exhaustion within eight hours, indicating that this effect is enhanced at ZT21 (Figure 1H). Blood glucose levels were reduced as expected after endurance exercise regardless of the time of day in both sedentary (Figure S1I) and active (Figure 1I) mice. Blood lactate levels were lower following endurance exercise at ZT21 than at ZT13 (Figure 1J). Note that the extreme running endurance of trained mice makes robust interpretation of differences between groups challenging because so many of the mice were able to run longer than the 8-hour maximum running time used in this study. Mice with higher lactate levels tended to be those that had reached the point of exhaustion consistent with the idea that lactate buildup is associated with exhaustion in endurance exercise.

### 3.3 RW activity only mildly impacts the core clock in skeletal muscle

To determine whether increased daily voluntary activity altered daily rhythms of gene expression in muscles, we established a new group of male c57BL/6J mice that were transferred to singly housed cages at the age of 9 weeks. Half were provided access to running wheels (RW) and half were not (sedentary, SED). Six weeks later, we collected muscles every four hours from each group over a 24-hour period (Figure 2A). We first used quantitative reverse transcriptase PCR (qPCR) to measure the basal expression of key circadian transcripts in quadriceps muscles. This analysis revealed that enhanced daily activity increases the daily amplitude of *Per2* expression but did not generally enhance the amplitude of core clock genes detected by this method (Figure 2B). While the profile of the PPARδ target gene *Pdk4* (Figure S2A) recapitulated our earlier finding of increased expression early in the day [17], it was not altered in response to voluntary running; nor was expression of the PPARδ coactivator *Ppargc1a* impacted (Figure 2B). Together, these results suggest that basal activation of PPARδ does not play a major role in the response to low intensity training afforded by voluntary running wheel access, although chronic activation of PPARδ in muscles is well established to enhance exercise capacity [29–31], and PPARδ is more robustly activated by acute intense exercise in trained mice than it is in sedentary controls [17].

**Figure 2.**
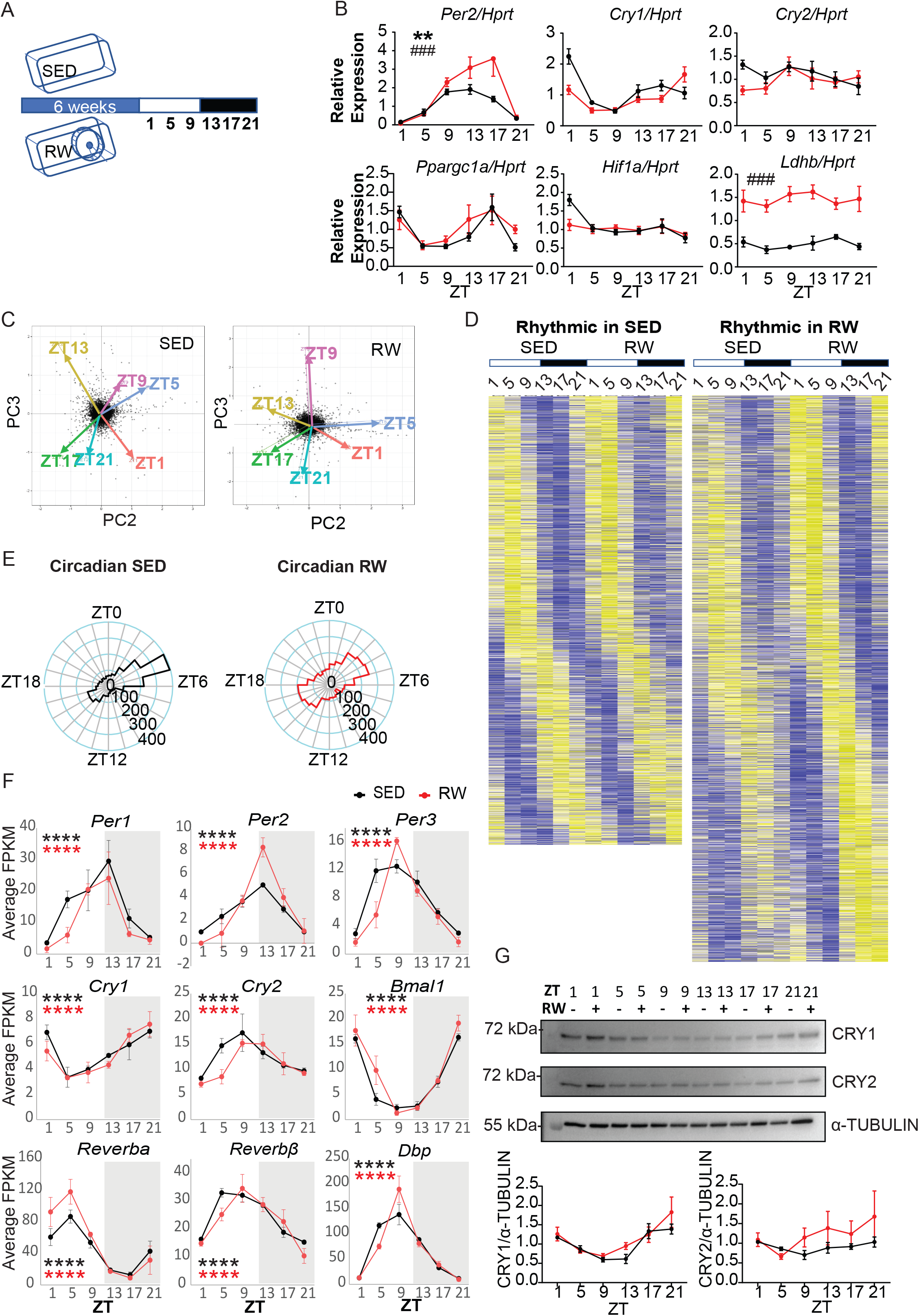
Daily exercise remodels the skeletal muscle circadian transcriptome. (A) Schematic diagram of the experimental setup. When C57BL/6J mice were 9 weeks old, they were transferred to individually housed cages without (SED) or with (RW) running wheels for six weeks. On the first day of week 7 (when they were 15 weeks old), muscles were collected every four hours starting at ZT1. (B) Expression of the indicated transcripts measured by qPCR in RNA isolated from quadriceps of sedentary (black) or voluntary running (red) male c57BL/6J mice collected at the indicated ZTs. Data represent the mean ± s.e.m. for 6 samples per condition, each measured in triplicate. ** P < 0.01, for a main effect of ZT, ### P < 0.001 of training effect by two-way ANOVA. (C) Principal component analysis reveals natural clustering of transcriptional phenotypes by ZT in both SED and RW datasets. (D) Heat maps representing expression levels (yellow high, blue low) of transcripts revealed by RAIN to be expressed with a circadian rhythm in plantaris muscles from sedentary (SED, left), or running wheel trained (RW, right) mice. Each horizontal line represents a single transcript. Transcripts are ordered by peak phase in SED (left) or RW (right) conditions. (E) Histogram showing distribution of peak phases for genes expressed rhythmically in plantaris muscles collected from sedentary (SED, black) or active (RW, red) mice. (F) Average detection of the indicated transcripts in sequenced RNA from sedentary (SED, black) or trained (RW, red) mice. **** P < 0.0001 for rhythmicity in sedentary (black) and trained (red) by RAIN. (G) Top, Detection of CRY1 and CRY2 by immunoblot in protein lysates prepared from plantaris muscles collected at the indicated ZTs. Each lane represents a unique plantaris sample. Data are a representative example of four sets of lysates from unique animals. Bottom, quantitation of Western blot data in which the signal for each sample was normalized to the average of all detected signals on the same Western blot exposure. Data represent the mean ± s.e.m. for four independent experiments.

Recently, several studies have demonstrated a close relationship between hypoxia signaling and circadian clocks, including suppression of HIF1α accumulation by the circadian repressors CRY1 and CRY2 [32; 33], HIF1α control of circadian clock gene expression [34; 35], and circadian modulation of *Hif1a* expression and of the transcriptional response to hypoxia [36; 37]. Training leads to a reduced transcriptional response to hypoxia [38]. Similar to the core circadian transcripts, we find that the mRNA expression of *Hif1a* was not significantly altered by running wheel activity (Figure 2B).

Increased expression of *Ldhb* has previously been recognized as a signature of the molecular response to low intensity training, and is believed to contribute to increased endurance exercise performance by enhancing the ability to utilize lactate as a fuel source [39; 40]. We measured robustly enhanced expression of *Ldhb* after chronic low intensity exercise training (Figure 2B). There was no impact of time of day on *Ldhb* expression in sedentary or trained mice. In sedentary animals, we measured greater *Ldha* expression in quadriceps during the late night and the reduction of its expression after training was greater at ZT17 than at other times measured (Figure S2A).

*Il-6* expression in muscles and circulating levels of IL-6 increase rapidly in response to intense exercise [41]. Although circulating levels of IL-6 have been reported to change in response to regular exercise in humans, the impact of training on IL-6 production remains unclear [42]. Unlike *Ldhb*, we did not measure any increase in *Il-6* expression after six weeks of voluntary running wheel training. Induction of *Il-6* is correlated with glycogen depletion. Low intensity voluntary running may not affect muscle glycogen content enough to increase *Il-6* expression relative to sedentary mice. Regardless, changes in *Il-6* expression within muscles do not seem to be required for the activity-induced enhancement of sprint exercise capacity at ZT21 in this study.

### 3.4 RW activity dramatically remodels daily rhythms of muscle gene expression

Because circadian clocks are best characterized as transcription-translation feedback loops and to take advantage of the rich literature on circadian regulation of transcription, we used high throughput RNA sequencing to measure the impact of daily voluntary wheel running on the daily profile of mRNA expression patterns in plantaris muscles to get a better sense of the impact of low intensity exercise training on rhythmic gene expression globally. We used plantaris muscles for this analysis because RNA prepared from whole plantaris muscles produces consistent transcriptome profiles across individual mice and similar global patterns of gene expression to those measured in quadriceps [43]. Principal component analysis provided confidence in the quality of our samples: The second (PC2) and third (PC3) principal components that emerge from an unbiased search for uncorrelated variables that maximize variance between samples reconstructs the circadian nature of the data in the correct time sequence (Figure 2C).

We used several complementary approaches to analyze genome-wide expression patterns. Using CircWave [44], robust harmonic regression, or RAIN [45], to analyze plantaris muscle RNA sequencing data from sedentary or active male mice, regardless of the false discovery rate, or amplitude or expression level filters employed, we find that voluntary running dramatically increases daily rhythms of gene expression within muscles (Figure 2D and Tables 1 and 2). For example, using the most stringent parameters to include only genes with the most robust expression and rhythmic amplitude reveals a striking 54% increase in the number of genes that are rhythmically expressed in plantaris muscles collected from mice housed with running wheels (1,446 genes) compared to those maintained in a sedentary state (938 genes) (Table 1). Furthermore, the phase distributions of rhythmically expressed genes are significantly different in active mice compared to sedentary mice (p < 0.001 by Watson’s two sample test for circular data, Figure 2E). This is also reflected in the outcome of detection of differential rhythmicity (DODR) analysis [46]: among the transcripts that are identified as rhythmically expressed in both sedentary and active mice using the stringent parameters in the previous example, a large fraction of them (680 transcripts) are found to exhibit altered rhythmicity between sedentary and active mice. However, as in quadriceps, this does not seem to reflect increased amplitude of core clock gene expression in plantaris (Figure 2F). Notably, while there is an 8-12 hour difference between the time of peak expression for *Cry1* mRNA and peak expression of *Cry2* mRNA (Figure 2F and Supplementary Tables S1 and S2), CRY1 and CRY2 protein rhythms in plantaris muscles are synchronous, peaking around the dark-to-light transition (Figure 2G).

**Table 1:** Circadian cutoffs and number of genes

| Expression cutoff | RAIN cutoff | Amplitude cutoff | n expressed | n circadian RW | n circadian SED |
| --- | --- | --- | --- | --- | --- |
| 0.1 | 0.05 | 0.10 | 15167 | 3155 | 2350 |
| 0.1 | 0.10 | 0.15 | 15167 | 1572 | 1010 |
| 1.0 | 0.05 | 0.10 | 11560 | 2824 | 2153 |
| 1.0 | 0.10 | 0.15 | 11560 | 1446 | 938 |

**Table 2:** DR cutoffs and number of genes

| Expression cutoff | RAIN cutoff | Amplitude cutoff | DODR cutoff | n DR | n DR, large amplitude change | N DR, amplitude larger RW | n DR, large phase change |
| --- | --- | --- | --- | --- | --- | --- | --- |
| 0.1 | 0.05 | 0.10 | 0.05 | 1212 | 835 | 682 | 324 |
| 0.1 | 0.05 | 0.10 | 0.10 | 2084 | 1278 | 1011 | 335 |
| 0.1 | 0.10 | 0.15 | 0.05 | 718 | 524 | 446 | 141 |
| 0.1 | 0.10 | 0.15 | 0.10 | 1120 | 720 | 607 | 145 |
| 1.0 | 0.05 | 0.10 | 0.05 | 1158 | 773 | 618 | 298 |
| 1.0 | 0.05 | 0.10 | 0.10 | 1992 | 1190 | 928 | 308 |
| 1.0 | 0.10 | 0.15 | 0.05 | 680 | 489 | 414 | 135 |
| 1.0 | 0.10 | 0.15 | 0.10 | 1061 | 675 | 564 | 139 |

To gain insight into processes that are coordinately influenced by circadian rhythms and activity, we used gene set enrichment analysis (GSEA) [47; 48] to identify pathways that are enriched among rhythmically expressed genes in plantaris muscles of sedentary or active mice. We generated lists of all expressed genes in plantaris muscles collected from either sedentary or active mice, ranked in order of the robustness of their rhythmicity as defined by the harmonic regression score (Supplementary Tables S3 and S4). Consistent with the similar rhythmic expression of core clock genes under both conditions, GSEA identified significant enrichment in both preranked lists for genes associated with circadian regulation of gene expression in the Gene Ontology [49; 50] Biological Processes subcollection of the Molecular Signatures Database (MSigDB) [51]. For genes that are rhythmically expressed in plantaris muscles of sedentary mice, GSEA found significant enrichment of multiple gene sets associated with RNA-mediated gene silencing, and of genes regulated by motifs matching several micro-RNA sequences (Supplementary Figure S2B). Notably, a search for enriched gene sets defined by the presence of any of 174 defined transcription factor binding motifs within 4 kb of noncoding sequence centered on their transcription start sites [51; 52] (TFT database) revealed that several nuclear hormone receptor (NR) binding elements are found in the vicinity of genes that are rhythmically expressed in plantaris muscles of active mice. Similar patterns of enhanced rhythmicity in plantaris muscles from active mice were also identified for target genes of hypoxia inducible factors (HIFs), and cyclic AMP responsive element binding protein (CREBP). Furthermore, these observations seem to reflect enhanced expression of transcripts in these gene sets during the dark phase when mice are most active (Supplementary Figures S2C-F).

To directly compare the rhythmic features of plantaris muscle transcriptomes from sedentary and active male mice, we employed DODR [46]. Consistent with the greater number of rhythmic transcripts detected in RNA from active mice described above, DODR revealed a large number of differentially rhythmic transcripts between the sedentary and active states (Table 2 and Supplementary Table S5). Notably, among the differentially rhythmic transcripts, there is a strong tendency for higher amplitudes of rhythmic expression in mice with voluntary running wheel access (Table 2). Depending on the cutoff values selected, between 135 and 335 transcripts exhibit a significant phase change in mice with access to a running wheel (Supplementary Table S2). Among these transcripts is a large group with peak expression early in the resting phase in sedentary animals (ZT0-ZT6) for which peak expression is advanced to the late active phase in mice housed with running wheels (ZT16-ZT24, Supplementary Figures S2G and S2H). Interestingly, analysis in gProfiler [53] reveals that this group of transcripts is highly enriched (p < 0.00001) for targets of RNF96 (a.k.a. TRIM28), which was recently shown to be activated by muscle contractions [54], consistent with the observed enhanced expression of these transcripts late in the active period in mice with access to running wheels.

We used differential expression analysis (DESeq2) to identify individual transcripts most impacted by the enhanced activity afforded by access to a running wheel (with zeitgeber time as a confounding factor) (Figure 3A and Supplementary Table S6). Among the 50 transcripts that are most up- or down-regulated in trained mice compared to sedentary mice, we found several well-established training-responsive transcripts as expected. The most overrepresented functional pathways associated with the upregulated transcripts include muscle structure development and morphogenesis; carbohydrate metabolism is overrepresented in the most downregulated genes as depicted by gProfiler and Revigo enrichment analysis (Figure 3B) [53; 55]. Myosin heavy chain isoform expression also changed as expected in response to low intensity exercise: the genes encoding myosin heavy chains 2 and 4 (*Myh2* and *Myh4*) were among the top increased and decreased transcripts, respectively (Figure 3C). This indicates that plantaris muscles contain a greater proportion of Type IIa fibers and reduced type IIb fibers after six weeks of voluntary running. This is consistent with the low intensity and long duration of the voluntary running activity that occurs with providing access to a running wheel, which is expected to enhance the proportion of oxidative fiber types within mixed muscle groups like the plantaris, and is expected to increase endurance exercise capacity.

**Figure 3.**
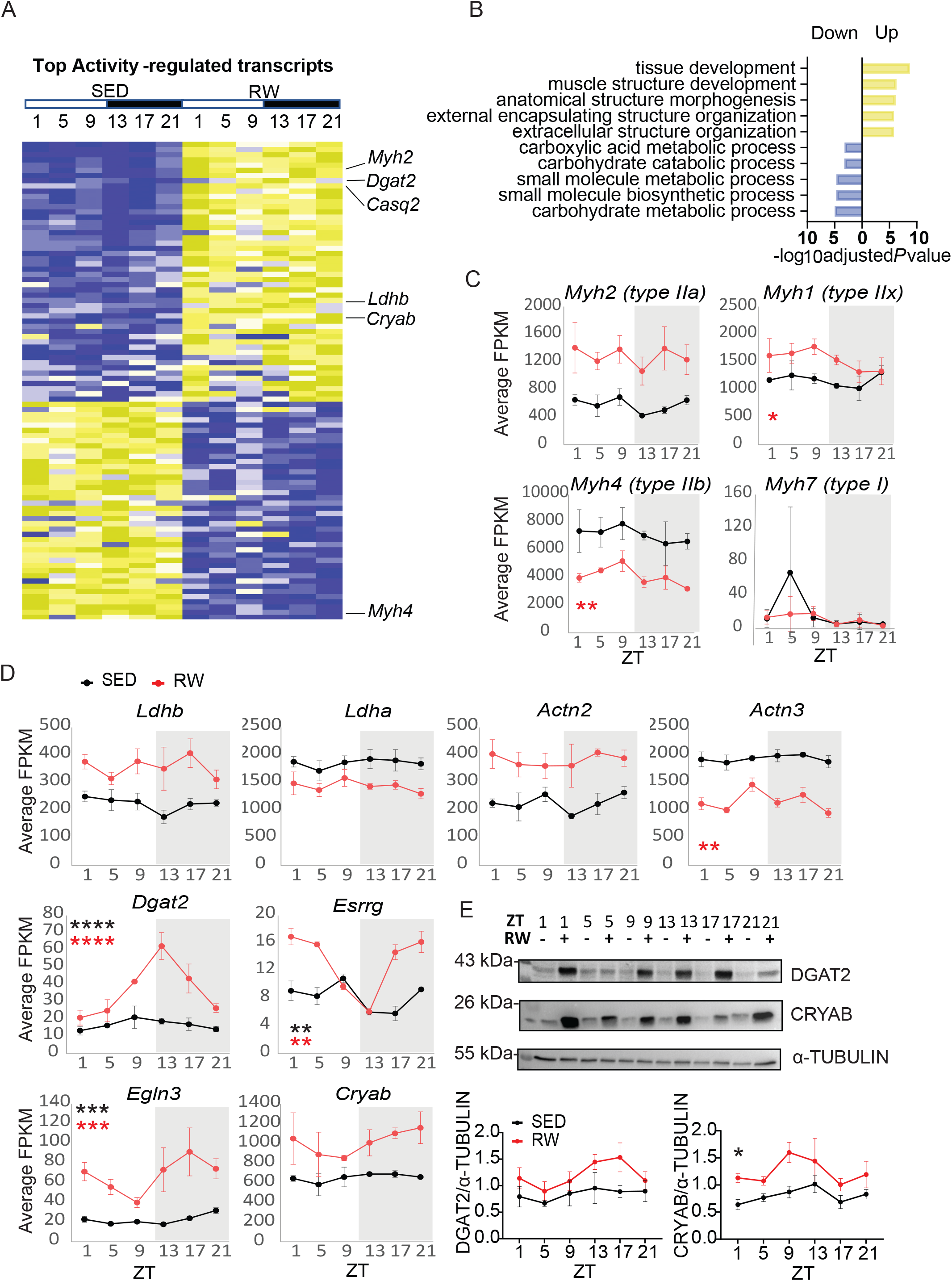
Metabolic and structural elements respond to training. (A) Heat map representing expression levels (yellow high, blue low) of the 100 transcripts revealed by DESeq2 to exhibit the greatest increase (top) or decrease (bottom) in transcription in response to training. Each horizontal line represents a single transcript. Individual gene names for select transcripts are indicated on the right. (B) Overrepresented functional pathways from (A) in gProfiler and simplified by Revigo analysis. (C,D) Average detection of the indicated transcripts in sequenced RNA from sedentary (SED, black) or trained (RW, red) mice. * P < 0.05, ** P < 0.01, *** P < 0.001 of rhythmicity in sedentary (black) and trained (red) by RAIN. (E) Detection of DGAT2 and CRYAB by immunoblot in protein lysates prepared from plantaris muscles collected at the indicated ZTs. Each lane represents a unique plantaris sample. Data are a representative example of four sets of lysates from unique animals. Bottom: Quantitation of Western blot data in which the signal for each sample was normalized to the average of all detected signals on the same Western blot exposure. Data represent the mean ± sem from four blots. * P < 0.05 for a main effect of daily exercise by two-way ANOVA.

As in quadriceps, *Ldhb* expression was consistently enhanced by activity in plantaris muscles (Figures 3A and 3D), supporting the idea that our protocol reproduces established signatures of enhanced endurance exercise capacity in response to training. *Ldha* expression was reduced in plantaris muscle of trained mice and was not significantly altered by zeitgeber time (Figure 3D). Together, these results suggest that lactate dehydrogenase isozyme expression is unlikely to contribute to the impact of time of day that we observed in sprint exercise capacity in trained mice.

In addition to the observed changes in myosin heavy chain expression and *Ldhb*, we found that transcripts encoding structural components of muscles and metabolic enzymes are overrepresented among those most responsive to increased daily activity. Intriguingly, several of these exhibit striking daytime-dependent expression, which may explain why their regulation by exercise state has not been observed previously (Figure 3D and Figure S3A). For example, the transcript encoding diacylglycerol O-acyltransferase 2 (DGAT2, an enzyme that catalyzes the final step in the synthesis of triglycerides) is slightly elevated in trained muscles during the day and its expression is six-fold higher in trained muscles compared to sedentary at ZT13 (Figure 3D). Furthermore, DGAT2 protein exhibits dramatically increased accumulation in plantaris muscles of trained mice only during the night, with peak abundance measured at ZT13-ZT17 (Figure 3E).

Similar to *Dgat2*, expression of the estrogen-related receptors beta (*Esrrb*) and gamma (*Esrrg*) was not strongly influenced by time of day in muscles from sedentary mice, consistent with prior studies of circadian regulation of nuclear hormone receptor expression in metabolic tissues [56]. However, their mRNA expression became dramatically dependent on time of day in plantaris muscles of mice housed with running wheels (Figure 3D and Figure S3A).

In addition, time of day affects the expression of transcripts that encode components of muscle cytoskeletal and sarcomere structures (Figure 3D and Figure S3A). The accumulation of the actin filament anchoring protein crystallin alpha B (CRYAB) is enhanced in plantaris muscles of active mice and seems to fluctuate over the course of the day (Figure 3E), but the impact of ZT on CRYAB protein accumulation is not statistically significant.

### 3.5 The impact of physical activity on the muscle transcriptome is transient

Importantly, the above analysis cannot distinguish between the immediate effect of recent physical activity and long-term transcriptional remodeling in response to exercise training. In order to gain insight into the acute and long-term impacts of daily activity on the muscle transcriptome, we measured the expression of selected transcripts that were robustly altered by physical activity in animals housed with running wheels for 10 weeks and then maintained in a cage with a running wheel that is either locked or free to move for up to 41 hours prior to collection of samples. This analysis revealed that termination of access to a freely moving running wheel rapidly alters gene expression (Figure S3B-C).

### 3.6 Voluntary RW activity and time of day coordinately influence glycogen metabolism

We used gene set enrichment analysis (GSEA, [48; 57]) with VST-normalized expression values for all detected transcripts as input to identify pathways defined in MSigDB [51; 58] that are impacted by daily physical activity. Transcripts involved in glycogen metabolism were enriched among those altered by access to a running wheel (Figure 4A-B). Specifically, the expression of several enzymes involved in glycogen breakdown is reduced in plantaris muscles of active compared to sedentary male mice (Figure 4C). In addition, the expression of several enzymes and regulatory factors that contribute to glycogen synthesis or breakdown depends on the time of day (Figures 4C and 4D). Glycogen synthase kinases 3 alpha (GSK3α) and beta (GSK3β) phosphorylate glycogen synthase and inhibit its activity. GSK3α and β are inactivated when they are phosphorylated (pGSK3α and pGSK3β). Despite their rhythmic mRNA expression, at the protein level we did not detect any difference in GSK3α or GSKβ between ZT13 and ZT21 (Figures 4E-F and Figure S4A-B). Hexokinase 2 (HK2) phosphorylates glucose to produce glucose-6-phosphate, which is required for entry into glycogen synthesis or glycolysis. Not only was *Hk2* transcript expressed rhythmically, HK2 protein was significantly lower in quadriceps muscles of active mice at ZT21 compared to those collected at ZT13 (Figures 4E and 4G).

**Figure 4.**
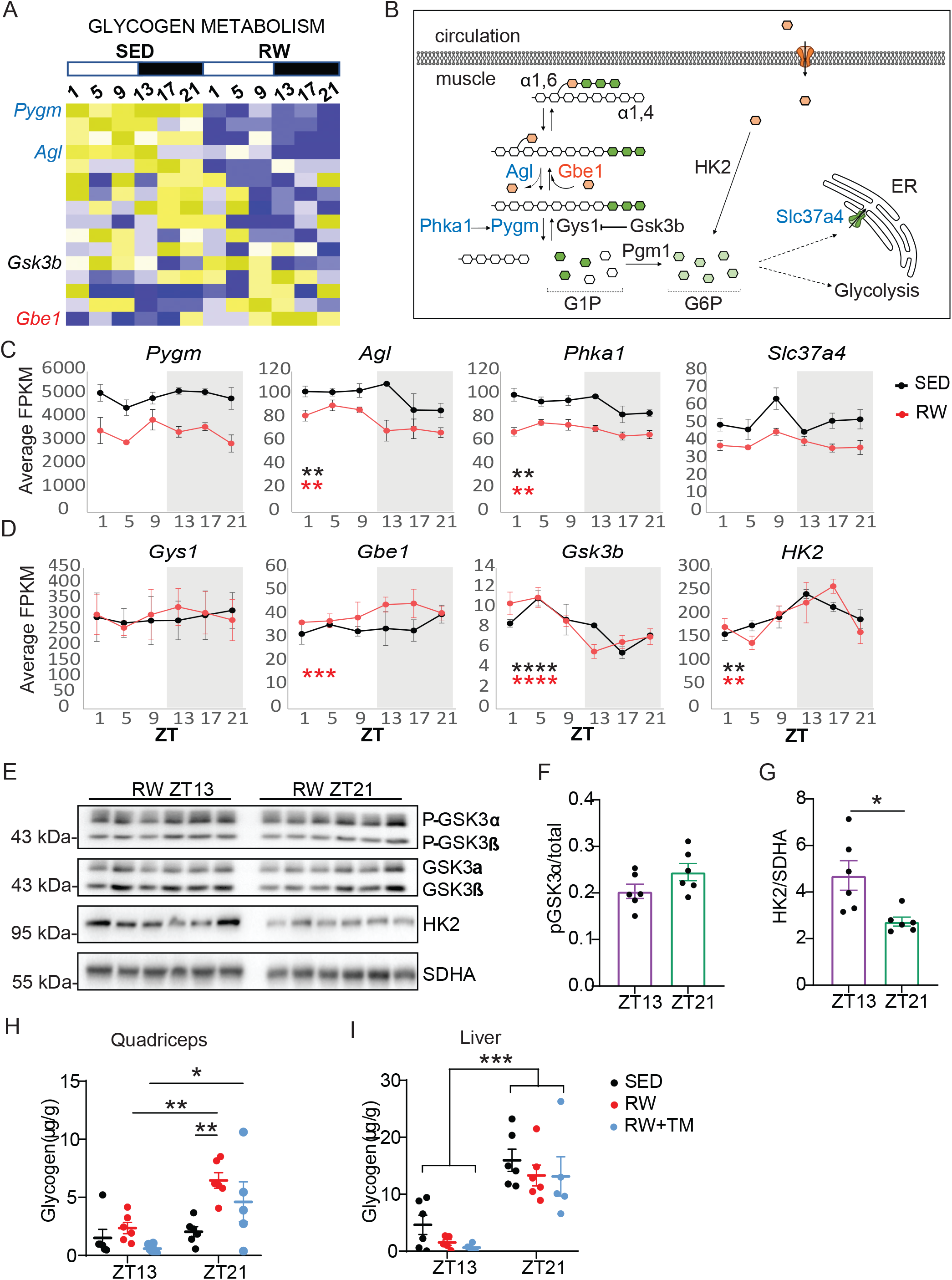
Daily exercise alters glycogen metabolism in a time of day dependent manner. (A) Heat map representing expression levels (yellow high, blue low) of transcripts involved in glycogen metabolism that are included in the “Mootha Glycogen Metabolism” gene set in MSigDB. Each horizontal line represents a single transcript. Individual gene names for select transcripts are indicated on the right. (B) Schematic diagram showing the function of selected enzymes in glycogen metabolism. Red and blue font indicate transcripts that are increased or decreased in plantaris muscles from mice housed with running wheels. (C, D) Average detection of the indicated transcripts in sequenced RNA from sedentary (SED, black) or trained (RW, red) mice. ** P < 0.01, *** P < 0.001, **** P < 0.0001 for rhythmicity in sedentary (black) and trained (red) by RAIN. (E) Detection of phosphorylated and total GSK3α and GSK3β, HK2, and SDHA by immunoblot in protein lysates prepared from plantaris muscles collected at the indicated ZTs. Each lane represents a unique plantaris sample. (F, G) Quantitation of the data shown in (E) * P < 0.05 by t-test. (H, I) Glycogen content measured in quadriceps (H) or liver (I) from sedentary (black) or trained (red) mice, or from trained mice that were subjected to exhaustive sprint exercise in the treadmill (green) at the indicated ZTs. * P < 0.05, ** P < 0.01, *** P < 0.001 by two-way ANOVA (factors: activity state, ZT) with post-hoc analysis.

Intramuscular glycogen provides fuel for energy production during intense exercise. The apparent regulation of glycogen synthesis and breakdown by both physical activity and time of day suggests that this could play an important role in the observed impact of time of day on maximum sprint speed in mice that have access to a running wheel for daily voluntary activity. Indeed, quadriceps muscles of male mice housed with running wheels contain three times more glycogen at ZT21 than they do at ZT13 (Figure 4H and Figure S4C). Liver glycogen content was higher at ZT21 in both sedentary and active trained mice (Figure 4I). While glycogen is almost entirely depleted at the time of treadmill failure in the mice tested at ZT13 as expected, a large amount of glycogen remains in both muscle and liver at ZT21 after the sprint exercise test (Figures 4H and 4I, RW+TM condition). Strikingly, while there is a tendency for intramuscular glycogen to be lower in mice that have undergone sprint testing (RW+TM, Figure 4H) at ZT21 it is not significantly different from that measured in similar mice that have not been run on a treadmill (RW, Figure 4H); liver glycogen is indistinguishable in livers collected at ZT21 with or without a treadmill sprint test (RW+TM vs RW, Figure 4I). These data indicate that glycogen utilization is reduced at ZT21 and that neither liver nor intramuscular glycogen are critical determinants of maximum running speed at ZT21 in mice after six weeks of voluntary exercise. Glycogen depletion is an established determinant of exhaustion during endurance exercise and lower glycogen concentrations in liver and muscle at ZT13 compared to ZT21 (Figure 4H and 4I) likely play a major role in the greatly reduced endurance that we (Figure 1H) and others [6] have measured at ZT13. While glycogen is not thought to play a major role in defining sprint performance, our data suggest that depletion of glycogen stores may unexpectedly contribute to limiting maximum running speed at ZT13. Together, these findings suggest that regulation of muscle glycogen metabolism is an important component of circadian modulation of exercise capacity in both sedentary and active mice. Furthermore, the reduced glycogen depletion at ZT21 and the large amount of glycogen remaining in muscle and liver after the treadmill sprint at ZT21 indicate that other pathways for energy production must be more readily available in mice that have had access to running wheels when they exercise at ZT21. Similar to their greater maximum speed, glycogen content of female quadriceps muscles was greater than that of males and while it also tended to be greater at ZT21 than at ZT13, the difference was not statistically significant (Figure S4D); liver glycogen content tended to be lower in female mice and also tended to be greater at ZT21 than at ZT13 regardless of activity (Figure S4E).

### 3.7 Activity impacts carbohydrate and lipid metabolism in a time of day dependent manner

Using the lists of all expressed transcripts ranked by harmonic regression scores as described above, we examined whether rhythmic expression of genes involved in other key metabolic pathways is influenced by daily exercise. This analysis revealed that transcripts involved in oxidative phosphorylation tend to have rhythmic expression patterns in sedentary animals but not in animals trained via voluntary wheel access (Figure 5A). Querying gene ontology [50] gene sets using GSEA revealed that transcripts involved in sodium ion transmembrane transport are enriched among those with greater rhythmicity in the trained state (Figure 5B). Of particular interest for exercise physiology, mRNAs encoding transporters that enable the import of the high energy metabolites creatine (SLC6A8), and inorganic phosphate (SLC20A1) exhibit elevated expression specifically late in the nighttime active period in trained plantaris muscles (Figures 5B and 5C).

**Figure 5.**
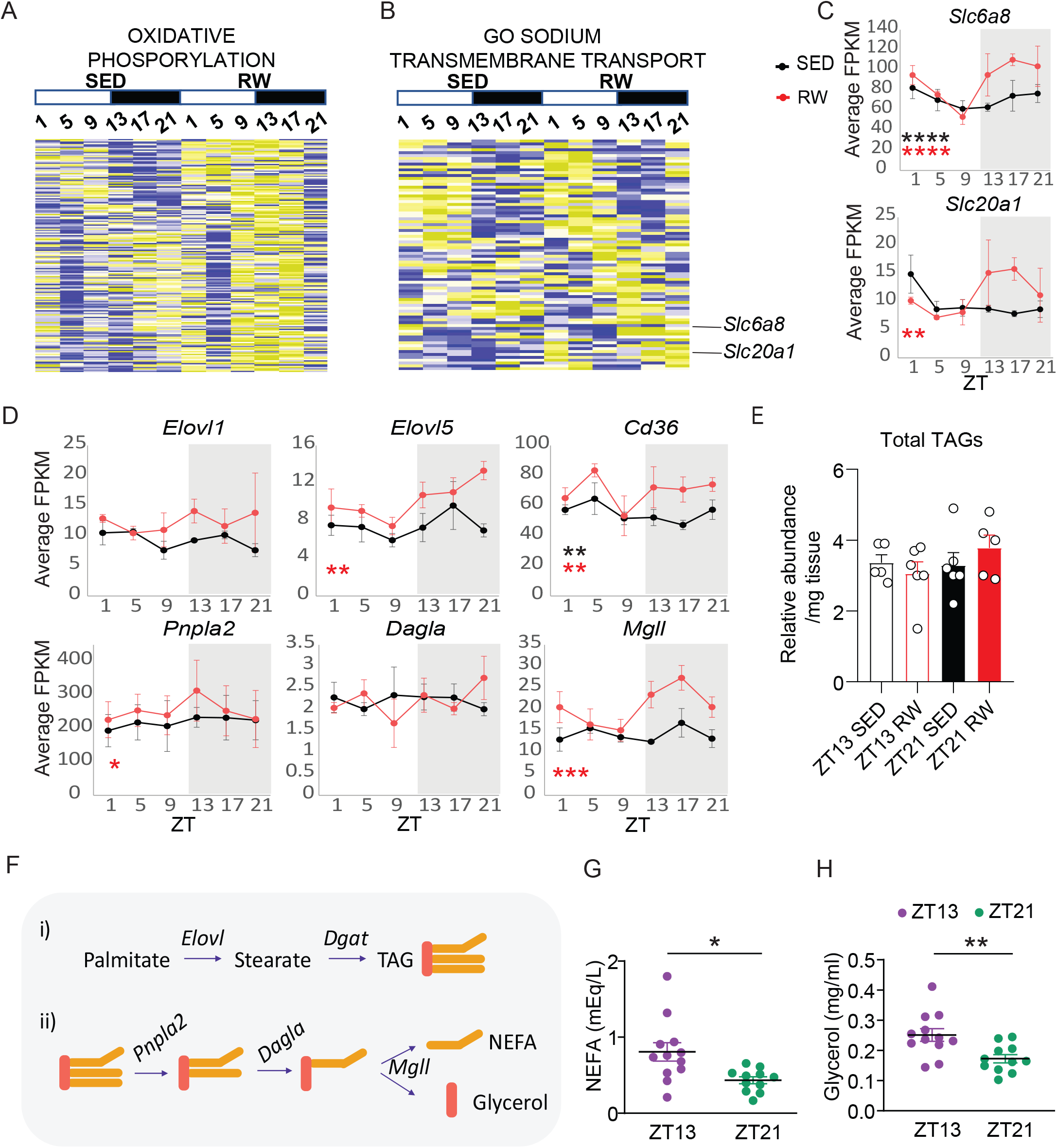
Training alters lipid metabolism in a time of day dependent manner. (A, B) Heat maps representing expression levels (yellow high, blue low) of transcripts included in the “Oxidative Phosphorylation” Hallmark gene set (A) or the “Sodium Transmembrane Transport” Gene Ontology gene set (B) in MSigDB. Each horizontal line represents a single transcript. (C) Average detection of *Slc6a8* and *Slc20a1*, (D) and lipid metabolic enzymes in plantaris muscle RNA sequencing. * P < 0.05, ** P < 0.01, *** P < 0.001, **** P < 0.0001 for rhythmicity in plantaris muscles collected from sedentary (black) and trained (red) mice by RAIN. (E) Total content of all triacylglyceride species measured by quantitative mass spectrometry in extracts from quadriceps muscles collected from sedentary (SED) or trained (RW) male mice at the indicated ZTs. Data represent the mean ± s.e.m. for six samples per condition. (F) Schematic diagram showing the function of selected enzymes in lipid metabolism. (G,H) Measurement of non-esterified fatty acids (NEFA) and glycerol in blood collected from mice housed with running wheels at ZT13 (purple) and ZT21 (green) * P < 0.05, ** P < 0.01 by two-way ANOVA.

The altered expression of transcripts and protein complexes involved in oxidative phosphorylation, and strong nighttime expression of lipid metabolic enzymes (Figure 5D), including DGAT2, for which we confirmed rhythmic expression at the protein level (Figure 3E), suggest that enhanced storage and utilization of lipids as fuel sources late in the active period of trained male mice could allow them to conserve intramuscular glycogen. Additionally, *Elovl1* and *Elovl5*, the only elongases detected above the threshold of 1 FPKM, seem to exhibit elevated expression in muscles from mice housed with running wheels (Figure 5D). We used targeted metabolomic profiling to measure the content of twenty-five fatty acid species with varying chain lengths and saturation. The total triacylglyceride (TAG) content of quadriceps muscles was affected neither by time of day nor by daily activity (Figure 5E). Among the 25 species of intramuscular lipids that we detected, there was a tendency for long-chain TAGs to be increased in quadriceps of active male mice collected at ZT21 but not at ZT13 (Figure S5A), consistent with the idea that voluntarily active male mice subjected to sprint exercise testing at ZT21 are able to conserve muscle glycogen stores by burning alternative fuel sources. Further supporting the hypothesis that active male mice increase both the synthesis and the utilization of intramuscular lipids at ZT21 compared to ZT13, they have reduced circulating non-esterified fatty acids (NEFA) and glycerol (Figure 5F-H) after sprinting to exhaustion at ZT21 compared to those exercised at ZT13. This is also consistent with increased expression of monoglyceride lipase (*Mgll*) at ZT13 compared to ZT21 that we measured in mice housed with running wheels (Figure 5D).

The major sources of energy production in exercising muscles are carbohydrates and lipids. We used targeted metabolite profiling to measure broad categories of metabolites in quadriceps muscles of sedentary and trained mice at ZT13 and ZT21 (Figure S5). The carbohydrate profile was significantly altered by exercise training in a time of day dependent manner. Glycolytic intermediates tended to be elevated at ZT21 compared to ZT13 in sedentary mice, and fumarate, malate and glutamate were significantly increased in active mice compared to sedentary mice at ZT13, but not at ZT21 (Figure S5C). Also, quadriceps contain more pyruvate and lactate at ZT21 than at ZT13 regardless of activity levels (Fig. S5C). Neither time of day nor activity level was associated with major changes in the profiles of lipids or amino acids that were detectable by this method.

## 4. Discussion

In contrast to a recent report [6], we did not observe any difference in sprint exercise capacity of sedentary male c57BL/6J when it was measured at ZT13 and ZT21. While research has not previously addressed the potential confounding impact of the use of electrical stimulation as an aversive stimulus to motivate treadmill running in rodents, another recent study used a manual method similar to the one that we employed to avoid the use of electrical stimulation in research using animals treated with chemotherapy [59]. Notably, they report similar maximum speeds in control animals as those that we observe in sedentary mice, further supporting the reproducibility of findings using this manual method. Further investigation will be required to understand the mechanistic basis for earlier termination of sprint exercise testing when an aversive electrical stimulus is used to motivate continued running. Regardless, it is important to recognize that electrical stimulation confounds exercise outcomes given the widespread use of this method to motivate continued treadmill running in exercise research using rodents.

In contrast to our finding that maximum sprint speed is indistinguishable between ZT13 and ZT21 in sedentary animals, male mice robustly increased their maximum sprint capacity in a striking time of day dependent manner after six weeks of running wheel training. As we [33] and others [60] have seen in prior studies, female mice tend to be more active than males; in addition, both maximum speed and tissue glycogen content are significantly greater in active c57BL/6J female mice than they are in males and there was no significant difference in these outcomes between female mice tested at ZT13 and those tested at ZT21. The explanation for this robust sexual dimorphism is unclear [60]. Differences in fatigability between males and females can be attributed to different muscle contractile and metabolic properties. For example, females oxidize more fat and less carbohydrate and amino acids than males do [61], and enhanced fatty acid oxidation is generally associated with greater endurance [62]. This may be related to the greater proportional area of skeletal muscle that is made up of type I fibers in females, which have smaller type II fibers than males [63] allowing better skeletal muscle fatigue resistance to endurance exercise. Moreover, differences in contractile properties such as lower rates of relaxation may contribute to more skeletal muscle fatigue resistance during sprint exercise. Type I fibers have a lower maximum sarcoplasmic reticulum Ca2^+^-ATPase activity and, therefore, female skeletal muscle metabolism is more suited to resynthesizing ATP from oxidative phosphorylation during exercise [64]. Notably, another study recently reported that female mice have greater exercise capacity at ZT22 than they have at ZT14, using a different exercise test that preferentially measures endurance exercise capacity as revealed by depletion of circulating glucose [65].

The effect of time of day on the beneficial impact of low intensity daily training on maximum sprint speed and exercise endurance in male mice enables us to identify molecular changes in response to daily activity that are associated with increased maximum running speed. Many established molecular responses to low intensity training [66] were equivalent at ZT13 and ZT21 while maximum speed was increased only at ZT21. Molecular differences between sedentary and active mice across the daily cycle indicates that activity-dependent remodeling of carbohydrate and lipid metabolism and metabolite transport within skeletal muscles underlie the observed time-dependent enhancement in performance.

We measured a dramatic remodeling of daily rhythms of gene expression in plantaris muscles in response to six weeks of voluntary wheel running. In general, a greater proportion of the genome exhibits daily rhythms of gene expression in mice performing daily voluntary exercise, consistent with a prior report that muscle groups containing a greater proportion of oxidative fibers exhibit more rhythmic transcription [67]. Consistent with the idea that oxidative muscles have greater transcriptional flexibility, overall genomic methylation is greater in type I (glycolytic) fibers isolated from human muscle biopsies than in type 2a (oxidative) fibers [68]. Further research is needed to understand how methylation and rhythmic transcription are dynamically altered by physical activity. Among the most striking examples of a transcript activated by training specifically during the nighttime active phase is that encoding an enzyme that catalyzes a rate-limiting step in lipid biosynthesis, DGAT2. We confirmed that DGAT2 protein abundance is dramatically increased in the late active phase and this, combined with elevated expression of elongating enzymes (e.g. *Elovl1, Elovl5*), likely influences the tendency for long chain lipids to be increased in quadriceps muscles of active mice isolated at ZT21. Previous studies have established the BMAL1-associated complement of mouse liver [69] and muscle [10] chromatin. In both cases, *Dgat2, Elovl1*, and *Elovl5* were found to be targeted by BMAL1 at multiple sites. Thus, BMAL1 can directly activate these lipid synthesizing enzymes and may enable rhythmic regulation of their enhanced expression in response to wheel running. Furthermore, genes involved in carbohydrate and lipid metabolism were enriched among those altered by genetic deletion of BMAL1 in muscle [14]. However, not all BMAL1 targets were so clearly altered by activity in a time of day dependent manner. While we saw an increase in rhythmic gene expression overall, many core clock transcripts unexpectedly did not exhibit enhanced rhythmic amplitude in active mice (Figs. 2B and 2E). Surprisingly, these data indicate that daily voluntary activity does not have a major impact on the muscle circadian clock. Mice in the active group continually had access to running wheels including on the day of sacrifice, and blocking wheel access tended to rapidly diminish activity-induced expression changes for several transcripts. Together, these findings are consistent with recent reports that the transcriptional response to exercise depends on the time of day [6; 13; 25], and indicate that the time at which exercise is performed may influence how it impacts physiology.

The modulation of lipid metabolism in response to voluntary wheel running is of particular interest in light of the so-called “athletes’ paradox” recognizing that intra-myocellular lipids are elevated in endurance trained athletes to levels similar to those observed in diabetic patients [70], while athletes’ insulin sensitivity remains robust. Coordinated elevation of lipid storage and oxidation in trained muscles likely explains the lack of pathological consequences [71]. Circadian modulation of these pathways may enable temporal coordination of lipid storage and utilization in the trained state to optimize performance while preventing detrimental impacts on overall metabolic health. While we did not observe an impact of time of day on total muscle TAG content, more detailed analyses would be required to measure intra-myocellular lipid content and to determine whether it is under circadian control.

Here, we demonstrate that voluntary running wheel activity consolidated in the early nighttime active phase led to increased running endurance regardless of the time of day, but enhanced maximal running speed only later in the night. Additional investigation will be required to understand how circadian clocks interact with exercise training of various types to coordinate changes in gene expression and physiology that ultimately determine exercise capacity.

## 5. Conclusions

Collectively, these data indicate that voluntary wheel running dramatically remodels daily rhythms of muscle gene expression in mice, despite resilience of the muscle core circadian clock to the influence of physical activity. Enhanced rhythmic expression of key metabolic transcripts in turn supports daily fluctuations in exercise performance that are exacerbated in mice that are physically active compared to those that are sedentary. Furthermore, our parallel investigations of circadian rhythms in gene expression and exercise performance support the emerging consensus that substrate selection for ATP production in muscles is under circadian control.

## Abbreviations

ZT: zeitgeber time
GSEA: gene set enrichment analysis
DODR: detection of differential rhythmicity
BMAL1: brain and muscle ARNT-Like 1
CLOCK: circadian and locomotor output cycles kaput
RW: running wheel
TM: treadmill
SED: sedentary
NEFA: non-esterified fatty acids
TAG: triacylglycerol
TFT: transcription factor targeting
NR: nuclear hormone receptor
qPCR: quantitative reverse transcriptase polymerase chain reaction

## 6. Acknowledgments

This work was funded by NIH grants R01 DK112927 (to K.A.L. and C.M.M.), R01CA234245 (to C. M.M.), and DK057978 (to R.M.E.). R.M.E. is an investigator of the Howard Hughes Medical Institute at the Salk Institute and March of Dimes Chair in Molecular and Developmental Biology. We thank Simon Schenk, Sabine Jordan, Robert Farese, Tobias Walther, Geraldine Maier, and Christoph Handschin for helpful discussions, sharing of technical expertise, equipment, and reagents, and/or critical reading of the manuscript, and T. Thomas, Y. Slivers, and J. Valecko for administrative assistance.

## Data statement

RNA sequencing data reported in this paper have been deposited in the National Center for Biotechnology Information (NCBI) Sequence Read Archive (SRA) database, BioProject ID PRJNA639978. Other original research data are available upon request.

