## Supplementary material for "Daily running enhances molecular and physiological circadian rhythms in skeletal muscle": Figure S1

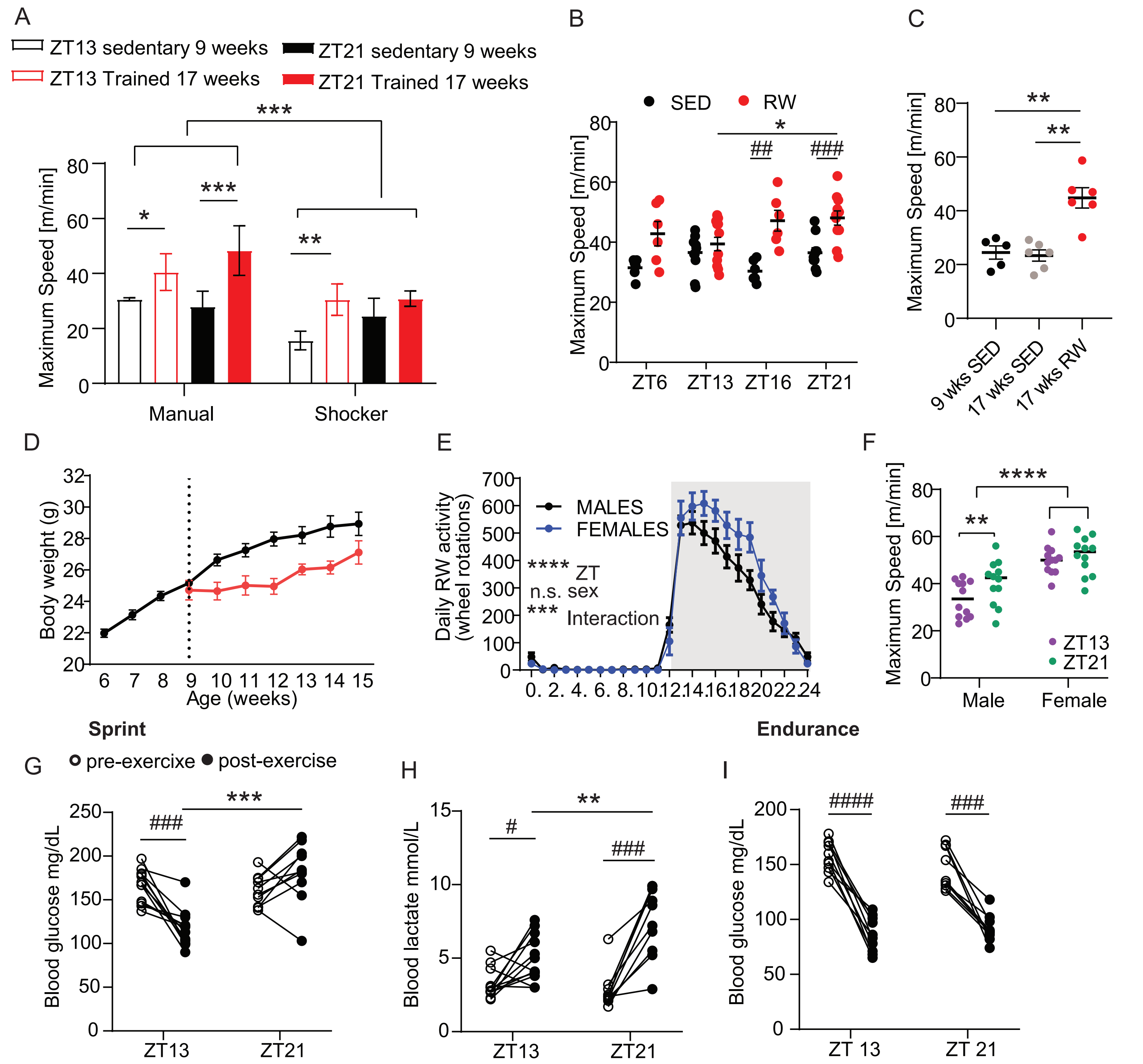


**Figure S1: Time of day impacts maximum speed in trained but not sedentary mice. Related to Figure 1.** (A) Maximum speeds achieved at ZT13 or ZT21 by c57BL/6J male mice using each of the systems shown in Figure 1A. Results of post-hoc testing are omitted for clarity. (B) Maximum speeds achieved by male C57BL/6J mice at the indicated zeitgeber times (ZT, hours after lights on before (black) or after (red) six weeks of voluntary access to running wheels. (C) Maximum speeds achieved by 9-week-old sedentary (black), 17-week-old sedentary (gray), or 17-week-old trained (red) male C57BL/6J male mice at ZT21. (D) Body weight of male mice from before the start of experiment (dashed line) and until end of experiment. (E) Temporal profile of running wheel activity of male (black) and female (blue) c57BL/6J mice during the period of voluntary access to running wheels. (F) Maximum speeds achieved by male and female C57BL/6J mice at ZT13 (purple) or ZT21 (green) after six weeks of voluntary access to running wheels. (G) Blood glucose and (H) lactate before (open black circle) and after (closed black circle) sprint test in sedentary male mice. (I) Blood glucose before and after endurance test in sedentary male mice. In (A, C, E, F) * $P < 0.05$, ** $P < 0.01$, *** $P < 0.001$, **** $P < 0.0001$ by two-way ANOVA followed by Tukey's multiple comparison test. In (B, G, H, I) each connected pair of open and filled circles represents measurements from the same individual mouse before (open circles) and after (closed circles) t he exercise test. * $P < 0.05$, ** $P < 0.01$, *** $P < 0.001$ by mixed-effects analysis with Sidak's multiple comparison test. # $P < 0.05$, ## $P < 0.01$, ### $P < 0.001$, #### $P < 0.0001$ by paired t-test.or paired t-test.
