## Supplementary material for "Daily running enhances molecular and physiological circadian rhythms in skeletal muscle": Figure S2

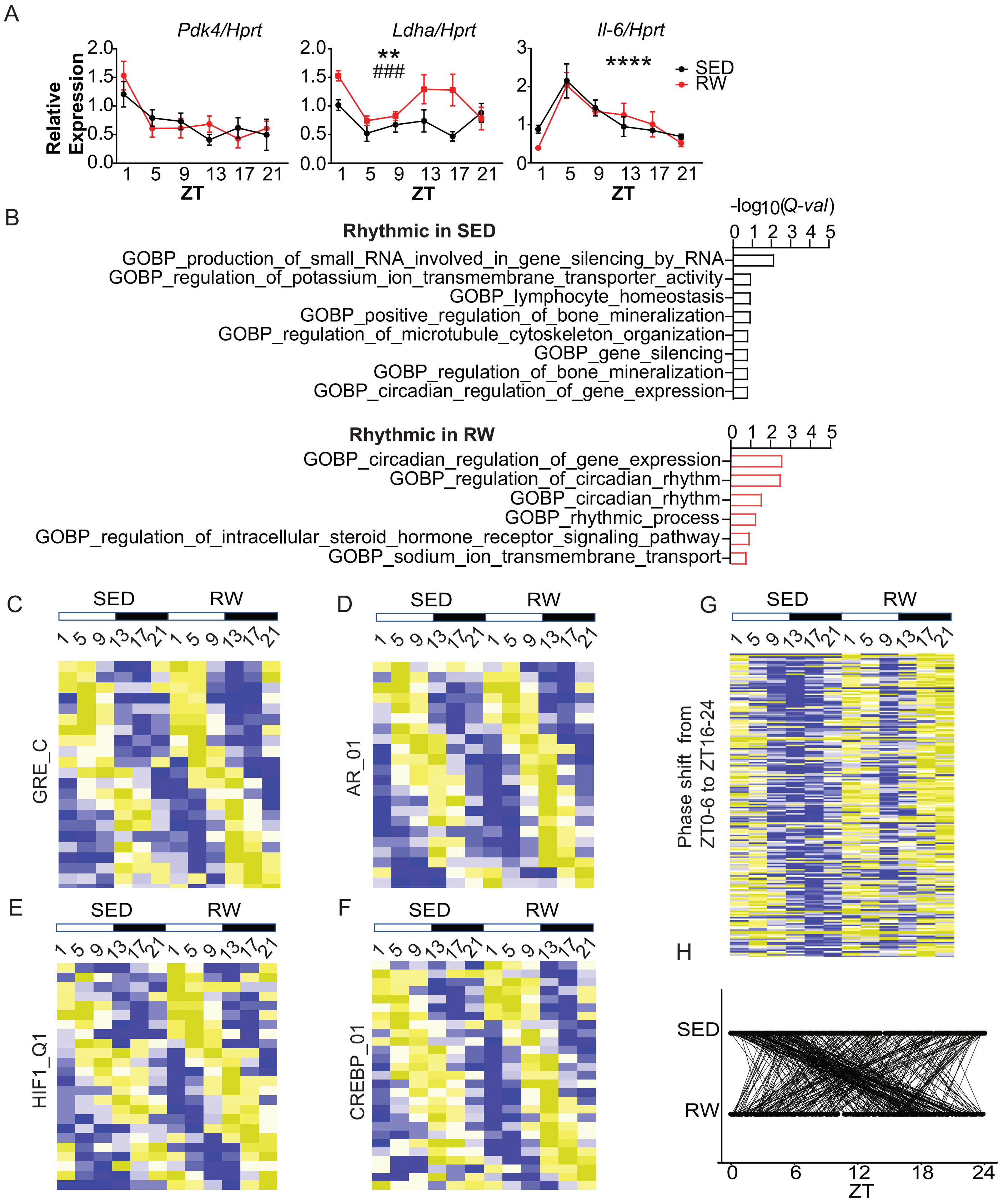


**Figure S2: Training remodels the skeletal muscle circadian transcriptome. Related to Figure 2.** (A) Expression of the indicated transcripts measured by qPCR in RNA isolated from quadriceps of sedentary (black) or trained (red) male c57BL/6J mice collected at the indicated ZTs. Data represent the mean ± s.e.m. for 6 samples per condition each measured in triplicate. ** $P < 0.01$, **** $P < 0.0001$ for a main effect of ZT, ### $P < 0.001$ of training effect by two-way ANOVA. (B) Overrepresented GOBP functional pathways from Figure 2D in gProfiler. (C-G) Heat maps representing expression levels (yellow high, blue low) for (C-F) targets of the glucocorticoid receptor (C), androgen receptor (D), hypoxia-inducible factor 1 (E), cyclic AMP responsive element binding protein (CREBP) (F), and for transcripts observed to undergo a phase shift from ZT0-6 in plantaris muscles collected from sedentary mice to ZT16-24 in plantaris samples collected from mice housed with running wheels (G). (H) A visualization of the phase changes from (G).
