## Supplementary material for "Daily running enhances molecular and physiological circadian rhythms in skeletal muscle": Figure S3

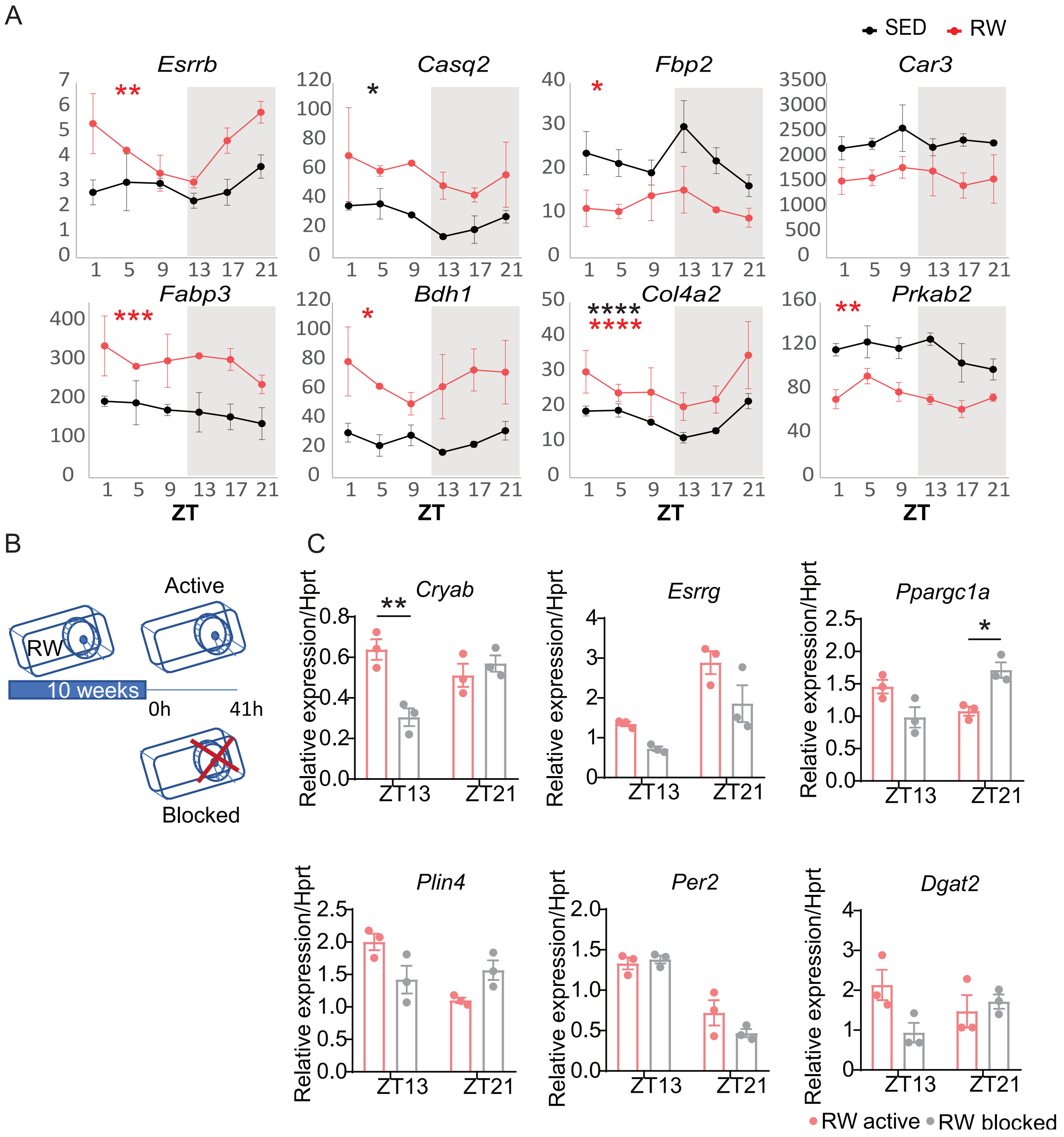


**Figure S3: Metabolic and structural elements response to training. Related to Figure 3.** (A) Average detection of the indicated transcripts in sequenced RNA from plantaris muscles isolated from sedentary (black) and active (red) mice at the indicated zeitgeber times (ZT, hours after lights on). Data represent mean ± s.e.m of VST-normilized read counts for 3 samples per condition. * P < 0.05, ** P < 0.01, *** P < 0.001, **** P < 0.0001 for detection of circadian expression in sedentary (black) and trained (red) by RAIN. (B) Schematic diagram of the experimental setup. c57BL/6J mice were placed into cages with running wheels for 10 weeks. Running wheel was either lock or free to move for up to 41h before dissection. (C) Expression of the indicated transcripts measured by qPCR in RNA isolated from plantaris of mice at ZT13 or ZT21 in an active (red) or locked (grey) running wheel. * P < 0.05, ** P < 0.01 by two-way ANOVA followed by Tukey's multiple comparison test.
