## Supplementary material for "Daily running enhances molecular and physiological circadian rhythms in skeletal muscle": Figure S4

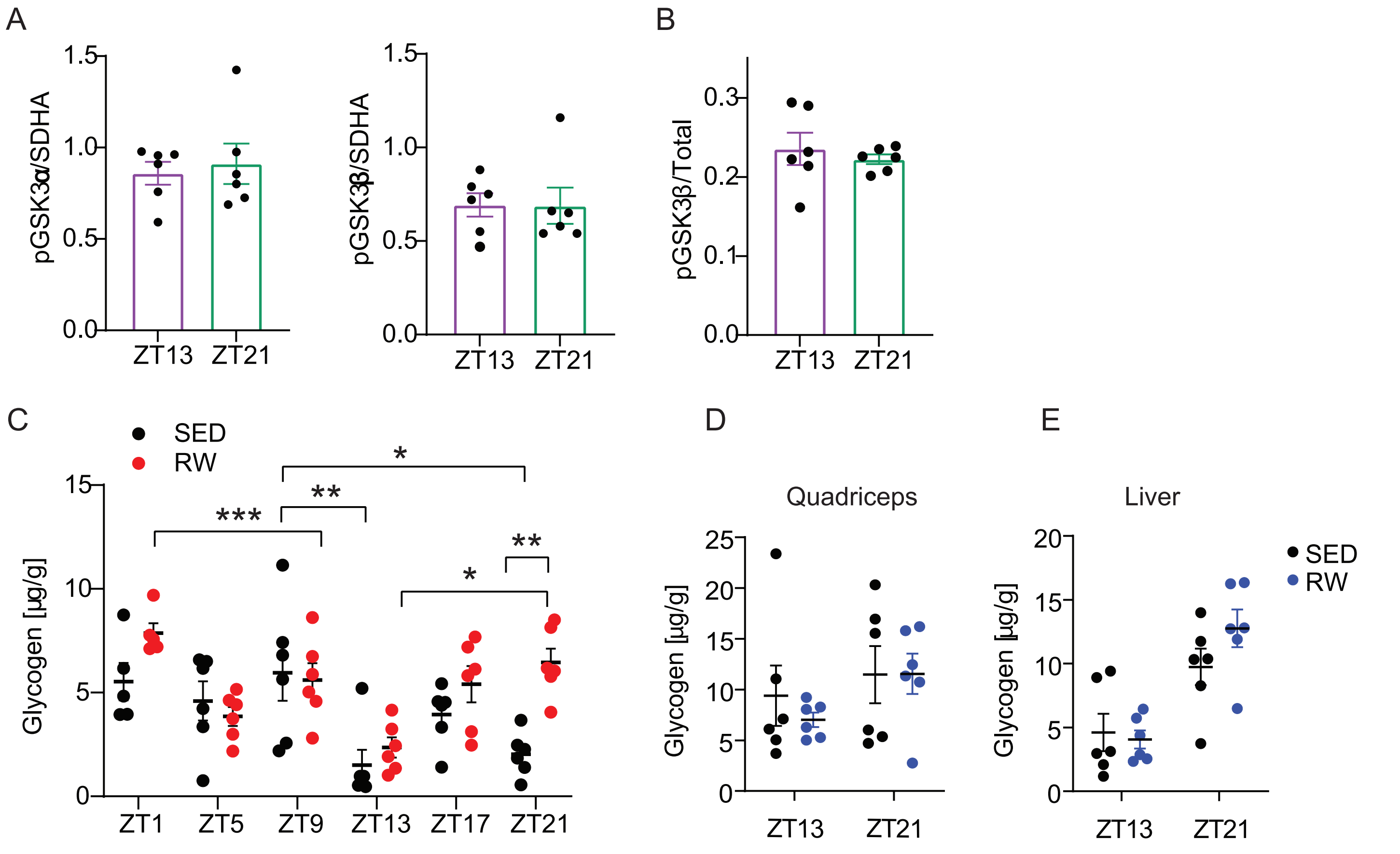


**Figure S4: Training alters glycogen metabolism in a time of day dependent manner. Related to Figure 4.** (A, B) Quantitation of pGSK3**α** and **β** data shown in Figure 4E. (C) Glycogen content measured in quadriceps from sedentary (black) or trained (red) male mice at the indicated ZTs. (D,E) Glycogen content measured in quadriceps (D) or liver (E) from sedentary (black) or trained (blue) female at the indicated ZTs. * $P < 0.05$, ** $P < 0.01$ by two-way ANOVA followed by Tukey's multiple comparison test.
