## Supplementary material for "Daily running enhances molecular and physiological circadian rhythms in skeletal muscle": Figure S5

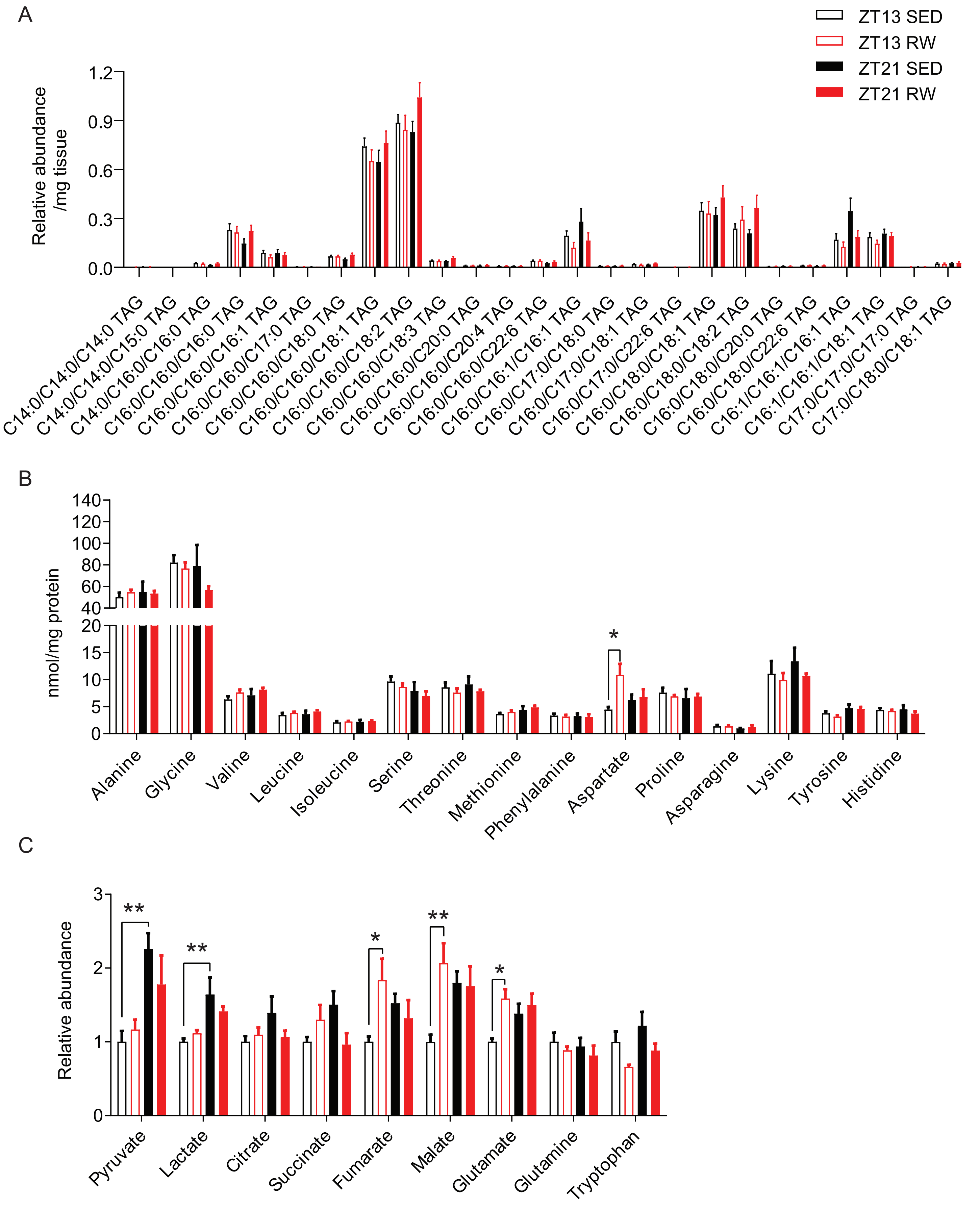


**Figure S5: : Training alters carbohydrate and lipid utilization in a time of day dependent manner. Related to Figure 5.** (A-C) Total content of individual triacylglyceride species (A), amino acids (B), and Krebs cycle intermediates (C) measured by quantitative mass spectrometry in extracts from quadriceps muscles collected from sedentary (black) or trained (red) mice at ZT13 (open bars) or ZT21 (filled bars). Data represent the mean ± s.e.m. of six samples per condition. * $P < 0.05$, ** $P < 0.01$ by ANOVA followed by Tukey's multiple comparison test.
